# Contribution of Uremia to *Ureaplasma*-Induced Hyperammonemia

**DOI:** 10.1101/2021.07.09.451868

**Authors:** Derek Fleming, Robin Patel

## Abstract

Lung transplant recipients (LTRs) are vulnerable to unexplained hyperammonemia syndrome (HS) in the early post-operative period, a condition typically unresponsive to non-antibiotic interventions. Recently, we showed that HS in LTRs is strongly correlated with *Ureaplasma* infection of the respiratory tract. It is not well-understood what makes LTRs preferentially susceptible to this phenomenon, compared to other immunocompromised hosts. *Ureaplasma* species harbor highly active ureases that convert urea to ammonia and CO_2_, utilizing the generated transmembrane potential to synthesize ATP. Post-operative LTRs commonly experience renal failure, resulting in uremia. We hypothesized that uremia could be a potentiating comorbidity to the development of HS secondary to *Ureaplasma* infection in LTRs by providing increased substrate for ureaplasmal ureases. We designed a novel dialyzed flow system to test the ammonia producing capacity of four isolates of *Ureaplasma parvum* and six isolates of *Ureaplasma urealyticum* in media formulations relating to normal and uremic host conditions. For all isolates, growth under uremic conditions resulted in significantly increased ammonia production over 24 hours, despite similar end-point bacterial quantities. Specifically, the isolates produced, on average, 1776.52 [standard deviation=263.98] μmol/L more ammonia when grown under uremic compared to normal conditions. This suggests that uremia, common in early post-operative LTRs, is a plausible contributing factor to the phenomenon of *Ureaplasma*-induced HS in this patient population.

## Introduction

Thousands of lung transplants are performed every year in the United States, with numbers anticipated to grow as availability and survivability continue to improve. A primary reason for greater survivability in recent years has been incorporation of strategies to minimize mortality caused by post-transplant infections. A contributor that has plagued lung transplant recipient (LTR) survival is hyperammonemia syndrome (HS), which occurs in around 4% of LTRs (1, 2). Ammonia (NH_3_) is a neurotoxin that readily transverses the blood-brain barrier and causes cerebral edema; chief mechanisms of NH_3_ neurotoxicity are illustrated in **Figure 1**.

**Figure 1.**
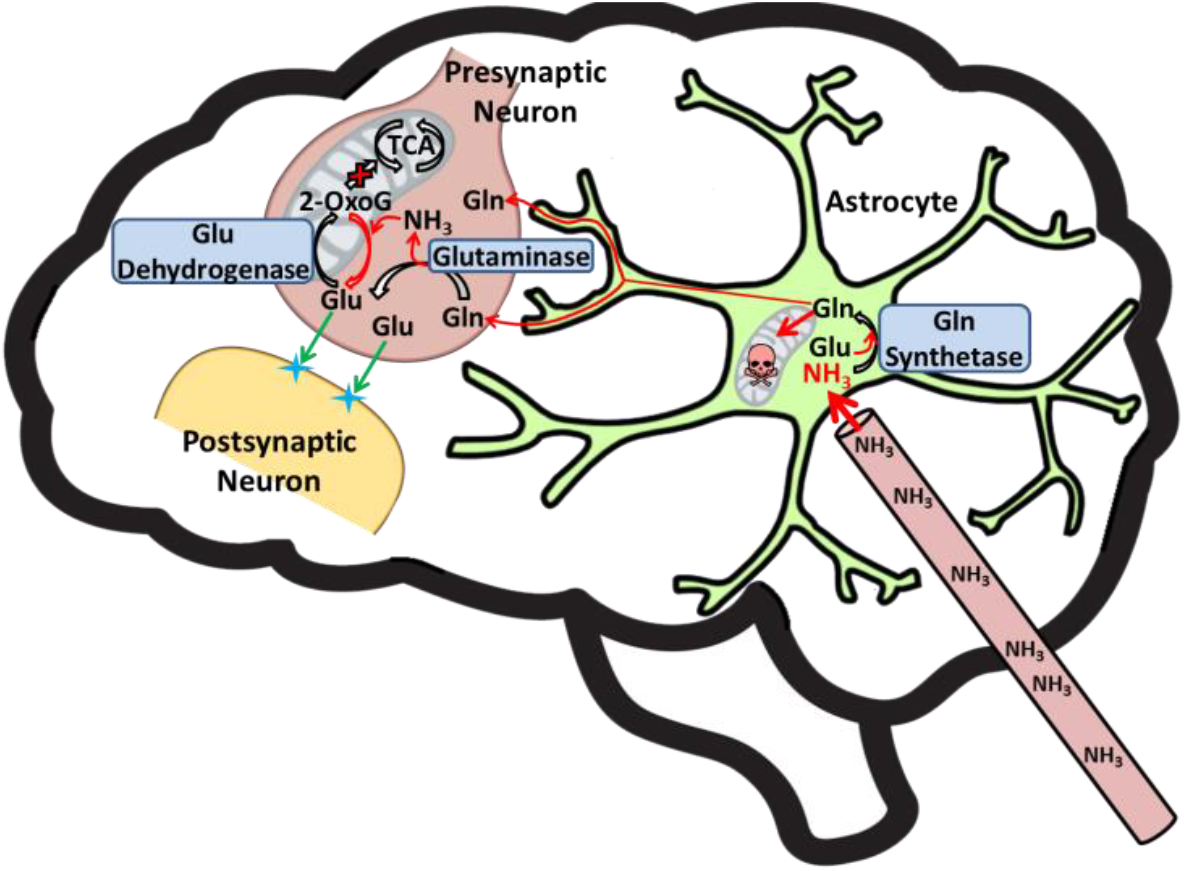
Ammonia (NH_3_) toxicity in the brain. Blood NH_3_ readily diffuses into astrocytes and other glial cells, combining with glutamate (Glu) to form glutamine (Gln) via glutamine synthetase. Some Gln is transported to neuronal cells, where it is converted into Glu via glutaminase, producing NH_3_ as a byproduct. Excess NH_3_ inhibits glutamate dehydrogenase, driving the reaction towards Glu, depleting 2-oxoglutarate (2-OxoG) and halting the tricarboxylic acid (TCA) cycle in both neurons and glial cells. Excess Gln in astrocytes is taken up by mitochondria, causing swelling and cell death.

HS following lung transplantation typically progresses from early identification of elevated blood NH_3_ levels or hyperammonemia (HA), as a cause of altered mental status resulting in confusion, lethargy, obtundation, and agitation, to eventual cerebral edema, resulting in seizure, coma, and often death (3–7). HS that presents in LTRs and, at lesser frequencies in other solid-organ transplant recipients (SOTRs), is atypical in that these patients do not have underlying liver disease or urea cycle disorders. Further, non-targeted interventional efforts to suppress endogenous NH3 production biochemically or physiologically and/or increase NH_3_ excretion have had minimal impact.

Recently, *U. urealyticum* and *U. parvum* were linked with HS in LTRs (8), where the airways of every LTR presenting with unexplained HS studied (n=13) tested positive for *Ureaplasma* species, likely of donor origin. Further evidence that *Ureaplasma* species are causative agents of HS in LTRs has come by way of *in vivo* studies with murine models, where it was shown that intratracheal and intraperitoneal infection with either *U. urealyticum* or *U. parvum resulted* in HA (9, 10). *Ureaplasma* species, which are normally considered commensal microbiota of the urogenital tract, produce a potent urease that splits urea into NH_3_ and CO_2_ as a means of ATP synthesis, powered by the generated NH_3_ gradient across the membrane (11, 12). Interestingly, 95% of ATP generated by *Ureaplasma* species is urea dependent, making it a requirement for growth (13). The high level of NH_3_ production from LTR *Ureaplasma* infection can exceed the capacity for detoxification by the host.

While unexplained HS has been described in non-LTR transplant patient populations (2, 14–28), the prevalence rate of ~4% seems highest, by far, in the LTR population. This suggests that the lung transplant scenario is specifically well-suited to microbial-driven HS. LTRs are also particularly vulnerable to uremia post-transplant (29, 30), potentially providing an abundance of substrate for ureaplasmal ureases, leading to NH_3_ overproduction, overwhelming the detoxification capacity of the host (**Figure 2**).

**Figure 2.**
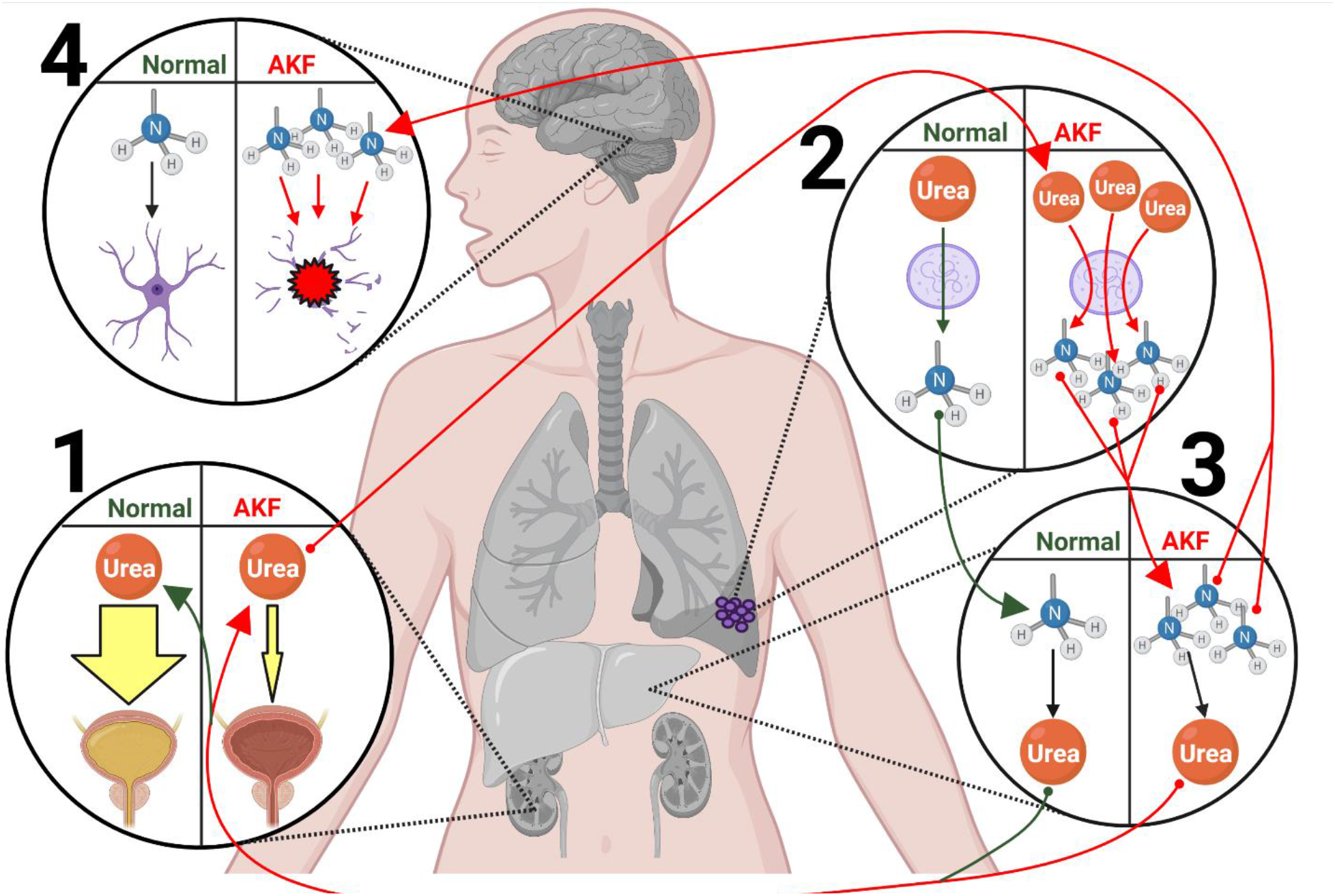
Hypothetical mechanism of *Ureaplasma*-induced hyperammonemia syndrome secondary to acute kidney failure in early post-operative lung transplant recipients. 1. Acute kidney failure (AKF) due to renal hypoperfusion results in decreased elimination of urea from the blood via urine excretion. 2. Increased blood urea availability provides additional substrate for ureaplasmal (purple cells) ureases in the infected respiratory tract, leading to greater ammonia (blue and grey NH_3_ molecule) production in comparison to normal blood urea concentrations. 3. Elevated serum ammonia levels overwhelm the detoxification capacity of liver urea cycle enzymes, leading to hyperammonemia. 4. Excess serum ammonia diffuses into astrocytes and other glial cells in the brain, causing them to swell and burst, resulting in cerebral edema and hyperammonemia syndrome. Created using BioRender.

Here, we investigated whether blood urea nitrogen (BUN) levels in LTRs could potentially contribute to production of pathological levels of NH_3_ by *Ureaplasma* species.

## Materials and Methods

### Study Isolates

The isolates of *U. parvum* and *U. urealyticum* investigated in this study are listed in **Table 1**. They include three respiratory isolates of *U. parvum* and five respiratory isolates of *U. urealyticum*, as well as one commercially available urogenital isolate of each species (from ATCC). Patient respiratory isolates are stored at the Mayo Clinic Infectious Disease Research Laboratory (IDRL). Isolates were grown to 10^7^ color changing units (CCU) using a *Ureaplasma* bioreactor, as previously described (31). 500 μL aliquots in U9 media (Hardy Diagnostics) buffered with 100 mM 2-ethanesulfonic acid (MES) at pH 6.0 were frozen at −80°C until use.

**Table 1.** *Ureaplasma* isolates studied.

| Species | Isolate # | Source |
| --- | --- | --- |
| *U. parvum* | IDRL-10774 | Bronchoalveolar Lavage Fluid |
| *U. parvum* | IDRL-11887 | Bronchoalveolar Lavage Fluid |
| *U. parvum* | IDRL-11264 | Sputum |
| *U. parvum* | ATCC-27815 | Urethritis |
| *U. urealyticum* | IDRL-10763 | Bronchial Washings |
| *U. urealyticum* | IDRL-10612 | Bronchoalveolar Lavage Fluid |
| *U. urealyticum* | IDRL-10611 | Bronchoalveolar Lavage Fluid |
| *U. urealyticum* | IDRL-11235 | Tracheal Secretions |
| *U. urealyticum* | IDRL-12698 | Bronchoalveolar Lavage Fluid |
| *U. urealyticum* | ATCC-27816 | Urethritis |

### Growth conditions

10 ml cultures of 10^5^ CCU/mL for each *Ureaplasma* isolate were encased in dialysis tubing (Specta/Por® Float-A-Lyzer®G2 1000kD Dialysis Device; G235073) and submerged in 250 ml of 100 mM MES-buffered U9, allowing measurement of NH_3_ levels over time. The entire device was incubated at 37°C, with fresh broth added and spent media removed, via flow at 2 ml/hour. Urea concentrations in the growth media were varied to mimic normal and high BUN levels (10 and 50 mg/dL, respectively) in the flow chamber and inflow. Samples were taken from cell-free portions of the flow chamber (outside the dialysis tube) and collected into microcentrifuge tubes at 0 and 24 hours, and NH_3_ concentrations tested using an NH_3_ Assay Kit (Abcam; ab102509). *Ureaplasma* cells in the dialysis tubing were quantified at each collection time.

## Results

For all *U. parvum* and *U. urealyticum* isolates tested, growth in media containing 50 mg/dl urea (uremic conditions) resulted in significantly greater NH_3_ production than in media containing 10 mg/dl urea (normal conditions) over 24 hours (**Figure 3**). Isolates grown under uremic conditions produced, on average, 1776.52 [standard deviation (SD)=263.98] μmol/L more NH_3_ than those grown in normal conditions. Differences in NH_3_ production between species was not significantly different, with *U. parvum* and *U. urealyticum* isolates producing an average of 1854.20 μmol/L (SD = 66.84) μmol/L and 1731.30 μmol/L (SD=131.23) more NH_3_, respectively, under uremic conditions. There was no noticeable difference between patient respiratory isolates and commercially available urogenital isolates. The average difference in NH_3_ production between uremic and normal conditions for all patient respiratory isolates was 1794.26 μmol/L (SD=291.40) *versus* 1800.95 μmol/L (SD=72.38) and 1655.44 μmol/L (SD=118.96) for the *U. parvum* (ATCC 27815) and *U. urealyticum* (ATCC 27816) urogenital isolates, respectively. 24-hour CCU counts between uremic and normal conditions was not significantly different (**Supplemental Figure 1**), indicating that NH_3_ production was not a result of different numbers of bacteria.

**Figure 3.**
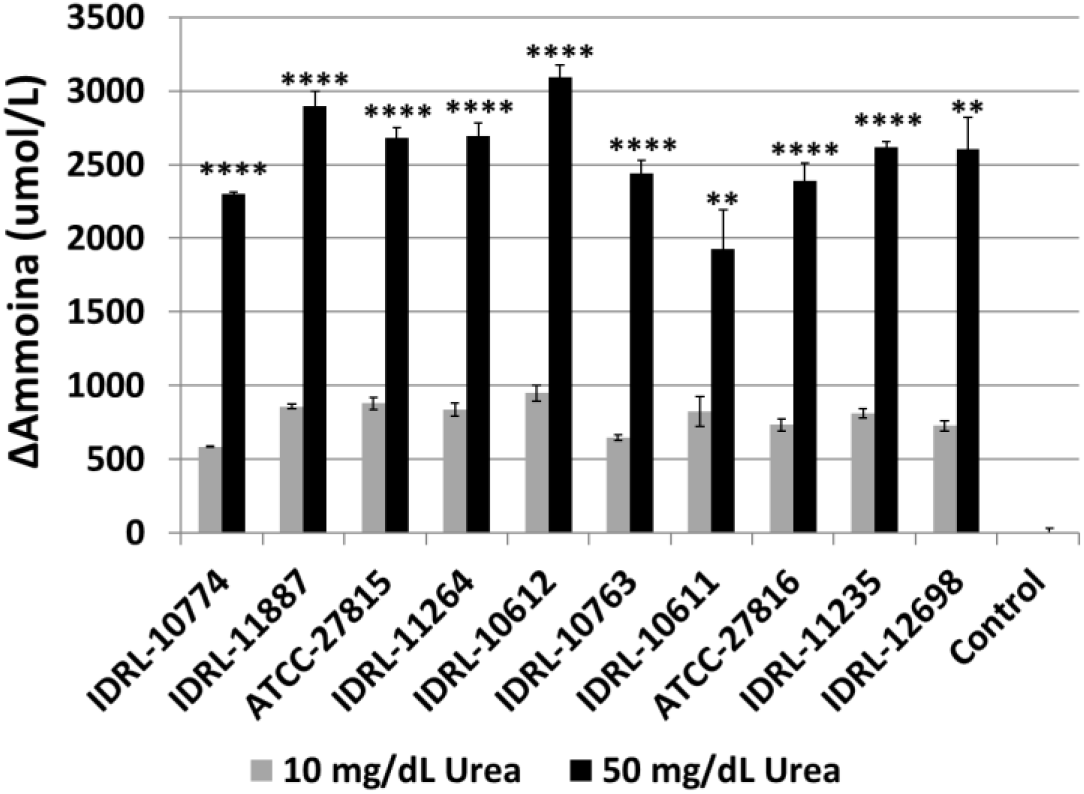
Average ammonia production by Ureaplasma isolates over 24 hours under normal and uremic conditions. Isolates of *U. parvum* or *U. urealyticum* were grown in the dialyzed flow system under normal (10 mg/dL urea) or uremic (50 mg/dL urea) conditions for 24 hours. N=3 per isolate per condition. Significance between conditions for each isolate was determined via two-tailed unpaired t-tests. **p ≤0.01, **p ≤0.0001.

## Discussion

Results of this study show that conditions representative of uremia resulted in elevated production of ammonia by all *Ureaplasma* isolates tested. The discovery that *Ureaplasma* respiratory infections were the cause of the previously unexplained phenomenon of non-hepatic HS in early post-operative LTRs (8) has led to an improvement in patient care and a reduction in mortality rates. Still, what makes LTRs particularly vulnerable to this phenomenon remains unknown. Here we investigated the potential impact of uremia as a product of acute kidney failure in the early postoperative period as a potentiating comorbidity. It has been estimated that as many as 75% of LTRs experience acute kidney failure post-operation due to renal hypoperfusion brought on by several factors, including 1) decreased circulating blood volume resulting from diuretic use to prevent pulmonary edema from leaky capillaries, 2) nephrotoxic effects of calcineurin inhibitors, and 3) reduced renal oxygenation due to post-operation hypoxia (29, 30, 32–34). We hypothesized that, among patients harboring a post-transplant *Ureaplasma* respiratory infection, elevated blood urea concentrations would provide greater substrate availability for ureaplasmal ureases, leading to the production of NH_3_ at sufficient levels to overwhelm the host detoxification capacity.

Using a novel dialyzed flow system, we measured the NH_3_ production of 10 isolates of *U. parvum* and *U. urealyticum* (8 clinical respiratory isolates and 2 commercially available urogenital isolates) over 24 hours under conditions representative of normal (10 mg/dl) and elevated (50 mg/dl) BUN. We found that, for all isolates tested, more ammonia was produced under uremic than normal conditions. Further, endpoint CCU counts were not significantly different for normal compared to uremic conditions, indicating that the greater ammonia production of uremic conditions can be credited to increased ureaplasmal urease activity. These results provide strong *in vitro* support for the hypothesis that post-transplant acute kidney failure is a contributing factor to *Ureaplasma*-induced HS in LTRs, serving as a reminder that the effects of specific comorbidities on disease of microbial metabolite production/overproduction should be more readily considered.

## Acknowledgements

The authors would like to thank Melissa Karau and Scott Cunningham for their assistance with this project. The authors would also like to thank the Clinical Microbiology Laboratory at Mayo Clinic, Rochester for providing the clinical isolates used in this study.

---

**Supplemental Figure 1.**
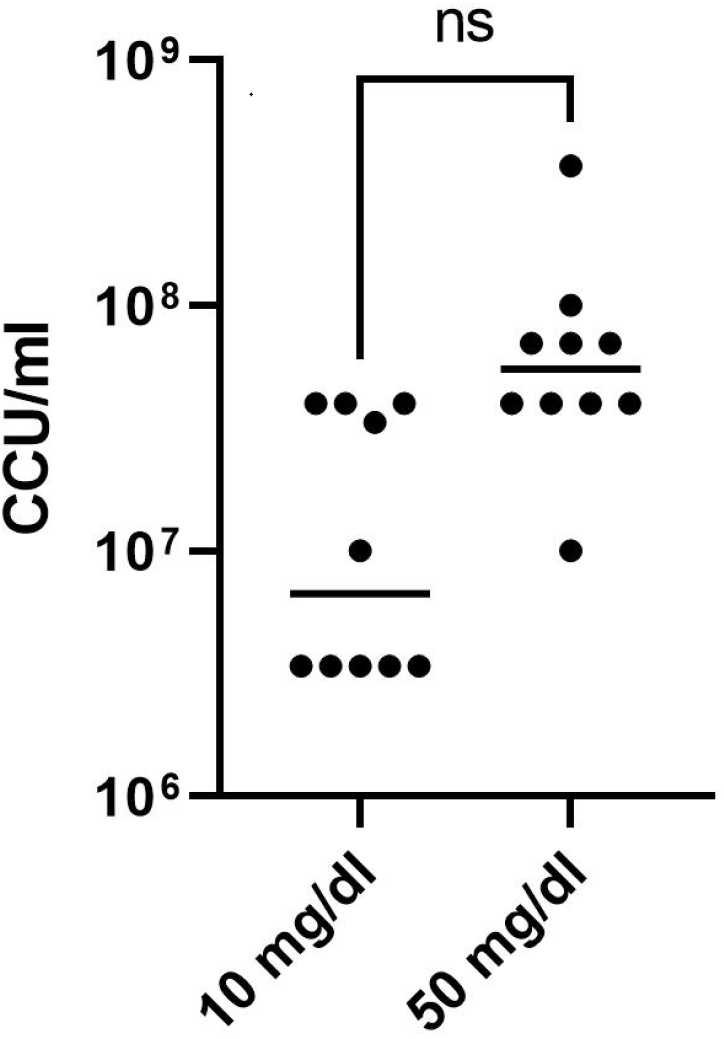
Differences in endpoint *Ureaplasma* quantities between normal and uremic conditions were insignificant. Isolates of *U. parvum* or *U. urealyticum* (N=3 per isolate per condition) were grown in the dialyzed flow system under normal (10 mg/dL urea) or uremic (50 mg/dL urea) conditions for 24 hours. Endpoint color changing units (CCUs) were quantified via serial dilution in 10B broth (Remel). Significance between conditions was determined via a two-tailed unpaired t-test.

## References

1. Lichtenstein GR, Yang YX, Nunes FA, Lewis JD, Tuchman M, Tino G, Kaiser LR, Palevsky HI, Kotloff RM, Furth EE, Bavaria JE, Stecker MM, Kaplan P, Berry GT. 2000. Fatal hyperammonemia after orthotopic lung transplantation. Ann Intern Med 132:283–7.

2. Krutsinger D, Pezzulo A, Blevins AE, Reed RM, Voigt MD, Eberlein M. 2017. Idiopathic hyperammonemia after solid organ transplantation: Primarily a lung problem? A single-center experience and systematic review. Clin Transplant 31.

3. Lichtenstein GR, Kaiser LR, Tuchman M, Palevsky HI, Kotloff RM, O’Brien CB, Furth EE, Raps EC, Berry GT. 1997. Fatal hyperammonemia following orthotopic lung transplantation. Gastroenterology 112:236–40.

4. Yoshida EM, Ostrow DN, Erb SR, Fradet G. 1997. Hyperammonemia after heart-lung transplantation. Gastroenterology 112:2162.

5. Rueda JF, Caldwell C, Brennan DC. 1998. Successful treatment of hyperammonemia after lung transplantation. Ann Intern Med 128:956–7.

6. Moffatt-Bruce SD, Pesavento T, Von Viger J, Nunley D, Pope-Harman A, Martin S, Ross P. 2008. Successful management of immunosuppression in a patient with severe hyperammonemia after lung transplantation. J Heart Lung Transplant 27:801–3.

7. Chen C, Bain KB, Iuppa JA, Yusen RD, Byers DE, Patterson GA, Trulock EP, Hachem RR, Witt CA. 2016. Hyperammonemia syndrome after lung transplantation: a single center experience. Transplantation 100:678–84.

8. Bharat A, Cunningham SA, Scott Budinger GR, Kreisel D, DeWet CJ, Gelman AE, Waites K, Crabb D, Xiao L, Bhorade S, Ambalavanan N, Dilling DF, Lowery EM, Astor T, Hachem R, Krupnick AS, DeCamp MM, Ison MG, Patel R. 2015. Disseminated *Ureaplasma* infection as a cause of fatal hyperammonemia in humans. Sci Transl Med 7:284re3.

9. Wang X, Greenwood-Quaintance KE, Karau MJ, Block DR, Mandrekar JN, Cunningham SA, Mallea JM, Patel R. 2017. *Ureaplasma parvum* causes hyperammonemia in a pharmacologically immunocompromised murine model. Eur J Clin Microbiol Infect Dis 36:517–522.

10. Wang X, Karau MJ, Greenwood-Quaintance KE, Block DR, Mandrekar JN, Cunningham SA, Patel R. 2016. *Ureaplasma urealyticum* Causes Hyperammonemia in an Experimental Immunocompromised Murine Model. PLoS One 11:e0161214.

11. Romano N, La Licata R, Russo Alesi D. 1986. Energy production in *Ureaplasma urealyticum*. Pediatr Infect Dis 5:S308–12.

12. Smith DG, Russell WC, Ingledew WJ, Thirkell D. 1993. Hydrolysis of urea by *Ureaplasma urealyticum* generates a transmembrane potential with resultant ATP synthesis. J Bacteriol 175:3253–8.

13. Dando SJ, Sweeney, Emma L., Knox, Christine L. 2019. *Ureaplasma*, p 1-28, Bergey’s Manual of Systematics of Archaea and Bacteria doi:10.1002/9781118960608.gbm01264.pub3.

14. Kiberenge RK, Lam H. 2015. Fatal hyperammonemia after repeat renal transplantation. J Clin Anesth 27:164–7.

15. Beeton ML. 2016. Possible missed diagnosis of *Ureaplasma* spp infection in a case of fatal hyperammonemia after repeat renal transplantation. J Clin Anesth 33:504–5.

16. Navaneethan U, Venkatesh PG. 2010. Idiopathic hyperammonemia in a patient with total pancreatectomy and islet cell transplantation. JOP 11:620–4.

17. Espinos J, Rifon J, Perez-Calvo J, Nieto Y. 2006. Idiopathic hyperammonemia following high-dose chemotherapy. Bone Marrow Transplant 37:899.

18. Mitchell RB, Wagner JE, Karp JE, Watson AJ, Brusilow SW, Przepiorka D, Storb R, Santos GW, Burke PJ, Saral R. 1988. Syndrome of idiopathic hyperammonemia after high-dose chemotherapy: review of nine cases. Am J Med 85:662–7.

19. Davies SM, Szabo E, Wagner JE, Ramsay NK, Weisdorf DJ. 1996. Idiopathic hyperammonemia: a frequently lethal complication of bone marrow transplantation. Bone Marrow Transplant 17:1119–25.

20. Sharp RA, Lang CC. 1987. Hyperammonaemic encephalopathy in chronic myelomonocytic leukaemia. Lancet 1:805.

21. Watson AJ, Chambers T, Karp JE, Risch VR, Walker WG, Brusilow SW. 1985. Transient idiopathic hyperammonaemia in adults. Lancet 2:1271–4.

22. Ho AY, Mijovic A, Pagliuca A, Mufti GJ. 1997. Idiopathic hyperammonaemia syndrome following allogeneic peripheral blood progenitor cell transplantation (allo-PBPCT). Bone marrow transplant 20:1007–8.

23. Frere P, Canivet JL, Gennigens C, Rebeix JP, Fillet G, Beguin Y. 2000. Hyperammonemia after high-dose chemotherapy and stem cell transplantation. Bone Marrow Transplant 26:343–5.

24. Tse N, Cederbaum S, Glaspy JA. 1991. Hyperammonemia following allogeneic bone marrow transplantation. Am J Hematol 38:140–1.

25. Uygun V, Karasu G, Daloglu H, Hazar V, Yesilipek A. 2015. Idiopathic hyperammonemia after hematopoietic stem cell transplantation: A case report. Pediatr Transplant 19:E104–5.

26. Snyder MJ, Bradford WD, Kishnani PS, Hale LP. 2003. Idiopathic hyperammonemia following an unrelated cord blood transplant for mucopolysaccharidosis I. Pediatr Dev Pathol 6:78–83.

27. Watanabe R, Okamoto S, Asahi A, Mori T, Takayama N, Ikeda Y. 2000. Idiopathic hyperammonemia following allogeneic bone marrow transplantation for refractory lymphoma. Jap J Clin Hematol 41:1285–8.

28. Placone N, Kao RL, Kempert P, Ruiz ME, Casillas JN, Okada M, Gibson JB, Maggi C, O’Brien K, Nattiv R, Gallant NM, Abrams SH. 2019. Hyperammonemia From *Ureaplasma* Infection in an Immunocompromised Child. J Pediatr Hematol Oncol doi:10.1097/MPH.0000000000001414.

29. Jacques F, El-Hamamsy I, Fortier A, Maltais S, Perrault LP, Liberman M, Noiseux N, Ferraro P. 2012. Acute renal failure following lung transplantation: risk factors, mortality, and long-term consequences. Eur J Cardiothorac Surg 41:193–9.

30. Lertjitbanjong P, Thongprayoon C, Cheungpasitporn W, O’Corragain OA, Srivali N, Bathini T, Watthanasuntorn K, Aeddula NR, Salim SA, Ungprasert P, Gillaspie EA, Wijarnpreecha K, Mao MA, Kaewput W. 2019. Acute Kidney Injury after Lung Transplantation: A Systematic Review and Meta-Analysis. J Clin Med 8:1713.

31. Fleming D, Karau M, Patel R. 2021. A novel bioreactor for the stable growth of *Ureaplasma parvum* and *Ureaplasma urealyticum*. J Microbiol Methods 181:106131.

32. George TJ, Arnaoutakis GJ, Beaty CA, Pipeling MR, Merlo CA, Conte JV, Shah AS. 2012. Acute kidney injury increases mortality after lung transplantation. Ann Thorac Surg 94:185–92.

33. Monnier A, Krummel T, Collange O, Haffner G, Hirschi S, Dégot T, Hannedouche T, Kessler R. 2015. Prevalence of scute and chronic renal failure after lung transplantation. J Heart Lung Transplant 34:S261.

34. Hanke N, Clajus C, Fegbeutel C, Walsemann D, Simon A, Haller H, Haverich A, Gueler F. 2012. Acute kidney injury after heart- and/or lung transplantation: Retrospective analysis of incidence, risk factors and outcome of 1200 patients: 2221. Transplantation 94.

